# Titration-WB: A methodology for accurate quantitative protein determination overcoming reproducibility errors

**DOI:** 10.1101/2024.07.16.603677

**Authors:** Alice Maestri, Ewa Ehrenborg, Olivera Werngren, Maria Olin, Carolina E. Hagberg, Matteo Pedrelli, Paolo Parini

## Abstract

Western blot (WB) technique is widely used to identify and quantify proteins in biological samples, but it is poorly accurate and precise for quantitative purposes. Merging the WB with the titration assay, we established a reliable method for protein quantification based on the slope of the linear function for signal-intensity/total protein loaded. Titration-WB unravels systematic and operator errors, compensates for problems in antibody affinity/avidity, and permits trustworthy inter-WB comparison.

## Introduction

Western blot (WB) is a widely used technique for the identification and quantification of proteins of interest in biological samples (Towbin, Staehelin, & Gordon, 1979). WB consists of five main steps: sample preparation, proteins separation by gel electrophoresis, proteins transfer onto a protein binding membrane, protein immunodetection using a specific antibody and signal detection and quantification. Due to the intrinsic nature of the biochemical and immunochemical reaction involved and due to the laborious and time-consuming steps, WB is a poorly accurate and precise technique when not employed properly (*e.g.* quantitative purposes). Several factors may affect the performance of WB, namely determination of the sample total protein concentration, identification of the appropriate lysis buffer, choice of the denaturing conditions, choice of electrophoresis running-time and of optimal gel matrix, appropriate protein binding-membrane, and optimization of the immunoreaction by identification of a specific antibody to quantify the protein of interest (Bass et al., 2017; Janes, 2015). Further, quantification of the WB often relies upon the normalization of the signal generated from protein of interest to the signal generated from a housekeeping protein. The latter is a major critical and limiting step of WB technique, where errors affecting the accuracy and precision can be easily introduced. Moreover, independently of the method of normalization, from the different housekeeping proteins to whole protein mass loaded, the obtained results from a single loaded sample protein are semi-quantitative and unreliable. Although many have proposed new ways to perform and quantify WB data (Ntoukas et al., 2021; Suzuki, Koura, Noguchi, Uchio-Yamada, & Matsuda, 2011; Taylor, Berkelman, Yadav, & Hammond, 2013), having accurate and precise quantitative measurements is still problematic. Here we describe how the application of the titration assay, originally developed to quantify vitamin C (McHenry & Graham, 1935), leads to accurate and precise quantitative WB results, and we name this method titration-Western Blot (t-WB). We have previously applied titration assay to WB for protein quantification in clinical and preclinical studies (Ahmed et al., 2019; Gnocchi et al., 2020; Minniti et al., 2020; Parini et al., 2008; Pedrelli et al., 2014; Pramfalk et al., 2022; Pramfalk et al., 2020). Here, we a aim to describe its framework, standardization, and validation.

## Results

The t-WB pipeline is described in Figure 1A. Firstly, each sample lysate should be prepared at three different serial dilutions, allowing the loading of three increasing protein masses of the same sample onto the gel. The dilution range needs to be carefully optimized so that immunodetection is possible for all dilutions, while aiming to an optimal signal to background ratio and to linearity, without reaching saturation. The use of at least three dilutions for each specimen is the first and fundamental improvement compared to the classical WB, in which each is usually tested by loading a single protein mass. Gel electrophoresis, transfer, and immunodetection follow the standard operating procedure optimised in each lab. From the three dilutions of each sample, three intensity signals are thus obtained. These intensity signals are then plotted against the corresponding protein mass loaded of the specimen. The resulting xy-graph is evaluated by the least squares method, since we operate under not saturating conditions, and the line of best fit for the regression is calculated (Figure 1A). The linear regression is also characterized by the coefficient of determination, or R-squared (R^2^), which is the statistical measure of how close the data are to the fitted regression line, estimating the percentage of variability explained by the linear function. In the t-WB the R^2^ is a key tool to assess the accuracy of sample dilution and sample loading. Evaluation of R^2^ also enables the understanding whether the most concentrated dilution points have reached saturation, although the linear relationship between sample concentration and band intensity should be warranted prior to the use of the t-WB analysis as also suggested by others (Taylor et al., 2013).

**Figure 1.**
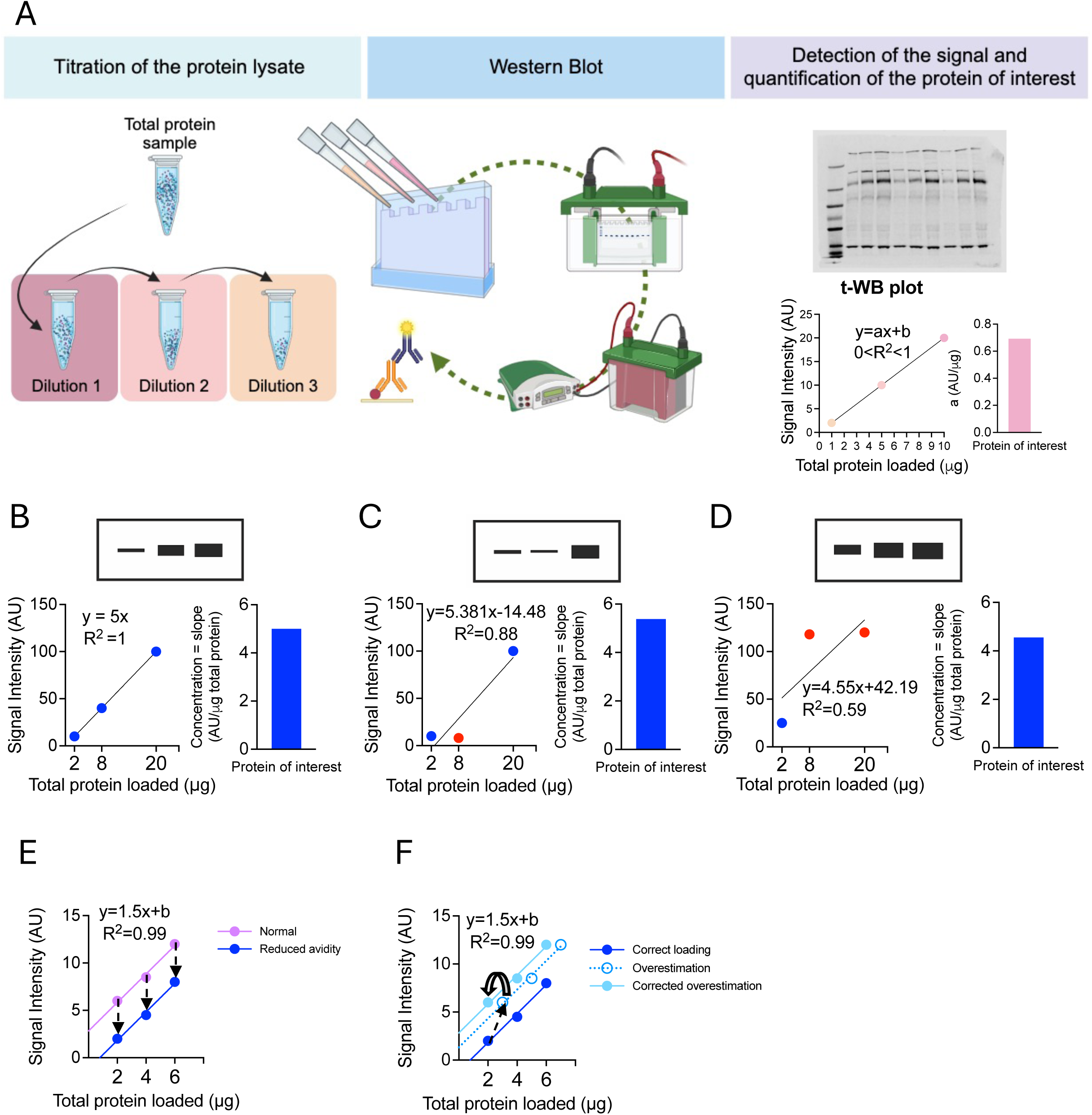
t-WB: pipeline and detection of immunoblotting errors. A) Schematic showing the t-WB pipeline. B) Schematic showing an ideal t-WB analysis: WB signal (upper insert) shows increasingly thick bands for increasing total protein mass loaded, for which the plotted signal results in a linear curve with a R^2^ of 1. The first derivative of the line of best fit is 5, thus the protein amount in this sample is 5 signal intensity units (AU)/total protein mass (μg), shown in the bar plot. C) t-WB is able to spot pipetting errors. The WB signal (upper insert) shows no increase in band intensity for the second data point, when plotted (in red) resulting in a low R^2^ value and reduced protein concentration (left bar plot); D) t-WB is able to detect saturation across sample dilutions (red points), again resulting in a reduced R^2^ value and protein concentration (left bar plot); E) t-WB corrects for reduced antibody affinity/avidity by a y-axis shift, without affecting the regression slope or protein quantification. F) t-WB corrects for overestimation of total loaded protein mass by a y-axis shift (dotted line), without affecting the regression slope or subsequent protein quantification.

The schematic in Figure 1B shows an ideal t-WB: the three sample dilutions result in three bands, which intensities are plotted against the correspondent total protein mass loaded. The resulting curve has an R^2^ of 1, which warrants no loading error nor saturation and, in this example a slope of 5 (Figure 1B; Supplementary Table 1A). However, in real world regression lines never pass the origin and the intercepts play a role in the assessment of the results, as discussed below. To obtain a quantitative measurement of the protein of interest, the first derivative *a* of the function *f*(*x*) = *ax*+*b,* describing the regression line, is calculated. It estimates the concentration of the target protein, expressed as arbitrary units of signal intensity *per* mass unit of total protein loaded (AU/mass). So, by calculating the first derivative it is possible to obtain a quantitative measurement of the concentration of the protein of interest, which for convenience is expressed in arbitrary unit. If the concentration of target protein expressed in mass or moles per total protein loaded is desired, *i.e.*, mass/mass or moles/mass, a standard curve with known concentration of the specific protein can be used. In our example in Figure 1B, *a* corresponds to 5 AU/mass. When running the WB in the classical way, the above-mentioned errors (*i.e.,* dilution, loading, and saturation) cannot be easily monitored or identified, and running a new WB blot will not necessarily help since the methodological approach will be the same and, in addition, the variability of immunodetection between blots would be an additional source of error. t-WB enables early detection of operator errors, such as inaccurate sample loading, by analysing the R² value, which decreases in proportion to the magnitude of errors present (Figure 1C; Supplementary Table 1B). Again, the R² can be easily used to monitor linearity, spotting saturation across the sample dilutions (Figure 1D; Supplementary Table 1C). The R^2^ will again decrease, and the slope *a* will be lower and inaccurate.

The mathematical function of the linear regression also allows the study of the intercept *b*, which is a measurement of the contribution of the background to the intensity measured in each point and informs on the signal to background ratio. The intercept *b* is also dependent upon the affinity/avidity of a primary antibody for the protein of interest, which is influenced by the co-localizing proteins present at the target site. Changes in affinity/avidity are reflected in shifts of the regression line along the y-axis. A reduction in affinity/avidity causes a reduced *b*, resulting from lower signal detection per unit of total protein loaded (Figure 1E; Supplementary Table 1D). Conversely, an increase in affinity/avidity leads to increased *b*. Further, discordant *b*-values between two sample replicates (*i.e.* two individual control samples run in two different WBs), may also reflect an erroneous quantification of the total protein concentration in one of the stock solutions prior to titration. Here, the regression lines shift on the x-axis either to the left, when the protein concentration is underestimated, or to the right, when overestimated (Figure 1F; Supplementary Table 1A). Since the operator is not aware of these potential sources of bias, running WB with a single amount of loaded specimen can give very inaccurate estimations of the concentration of the protein of interest, and standardization by using a loading control only partly helps. In t-WB, all the above-mentioned sources of bias do not interfere with the quantification of the protein of interest as the coefficient *a* (or slope) is not affected by these shifts, maintaining high accuracy in the determination of the concentration of the protein of interest.

To illustrate the accuracy of the t-WB method compared to the classical WB method, we evaluated the expression of the lipid droplet-associated protein perilipin 2 (PLIN2) in THP- 1 cells upon different lipid loading conditions (Figure 2; Supplementary Table 2). In t-WB, the signal intensities resulted from the three different dilutions of each condition were plotted against their respective masses of total protein loaded (Figure 2B; Supplementary Table 2B); the first derivative *f’(x)=a* was calculated and plotted to evaluate the effects of the different lipid loading treatments (Figure 2C; Supplementary Table 2B). Subjecting THP-1 cells to either 25 or 50 μg/mL of oxidized low-density lipoprotein (oxLDL) in the culture media led to about a 3.4-fold increase of PLIN2 protein, while adding 400 μM of bovine serum albumin (BSA)- coupled oleic acid (OA) increased the PLIN2 concentration 4.3-fold as compared to control cells. To compare these results with the classical WB approach, we also probed and quantified the signal of the housekeeping protein β-actin for each sample to correct for loading errors (Figure 2D-F; Supplementary Table 2C-D). Notably, we found a variable β-actin signal, particularly affected by OA incubation, which reduced its intensity (Figure 2F; Supplementary Table 2C-D). We could exclude that the above β-actin signal variability resulted from a loading error of the samples because linearity is present as revealed by the very high R^2^ (0.98) of the regression line obtained with t-WB for PLIN2 in the same specimen. In the classical WB methods, variance in the housekeeping protein in response to treatment is unfortunately a source of bias that is systematically neglected and hard to detect when only a single mass of total protein is loaded onto the gel. However, this source of bias is very relevant, and it dramatically affects the quantification of the protein of interest (in this case PLIN2), leading to faulty conclusion. Next, the changes in PLIN2 expression upon lipid treatment were quantified using the classical WB method, using only the point from the 1 μg of total protein mass loaded and adjusting for the respective β-actin signal (Figure 2I, first panel; Supplementary Table 2E). This normalisation resulted in a grave overestimation of the effect of OA treatment on PLIN2 expression (Figure 2G, first panel; 16.8- *vs*. 4.3-fold with t-WB; Supplementary Table 2E). Using the 10 μg of total protein mass loaded data point instead, once adjusted for the β-actin signal, leads to a more accurate quantification (5.7- *vs.* 16.8-fold; Figure 2G, first and third panels; Supplementary Table 2E). However, none of the loaded concentrations, including the 5 μg of total protein mass loaded data point, were able to correctly quantify changes in PLIN2 expression upon OA-treatment, exposing the limitations of accuracy when performing and quantifying WB data in the classical way (Supplementary Table 2E). This lack in accuracy is also evident when comparing the effect of treating THP-1 cells with two different concentrations of oxidized low-density lipoprotein (oxLDL). Quantification using the samples where 1 μg or at 5 μg of total protein mass was loaded and normalized to β-actin, suggested that PLIN2 protein expression increases in a dose-dependent manner upon oxLDL loading (Figure 2G, first and second panels; Supplementary Table 2E). However, this dose-dependent increase in PLIN2 protein expression disappeared when quantification and standardization by β-actin were done at 10 μg of total protein mass loaded data point (Figure 2G, third panel; Supplementary Table 2E). Using the t-WB we see that 25 or 50 μg/mL of oxLDL treatment induces a comparable fold increase of PLIN2 as compared to vehicle treated cells (Figure 2C: 3.4-fold both at 25 and 50 μg/mL of oxLDL respectively *vs.* control). In conclusion, the t-WB method allowed correct quantification of changes in PLIN2 protein expression upon different treatments quantitatively without relying to housekeeping proteins, avoiding this additional sources of bias while asserting operator errors influenced the results (Goasdoue, Awabdy, Bjorkman, & Miller, 2016; Li & Shen, 2013).

**Figure 2.**
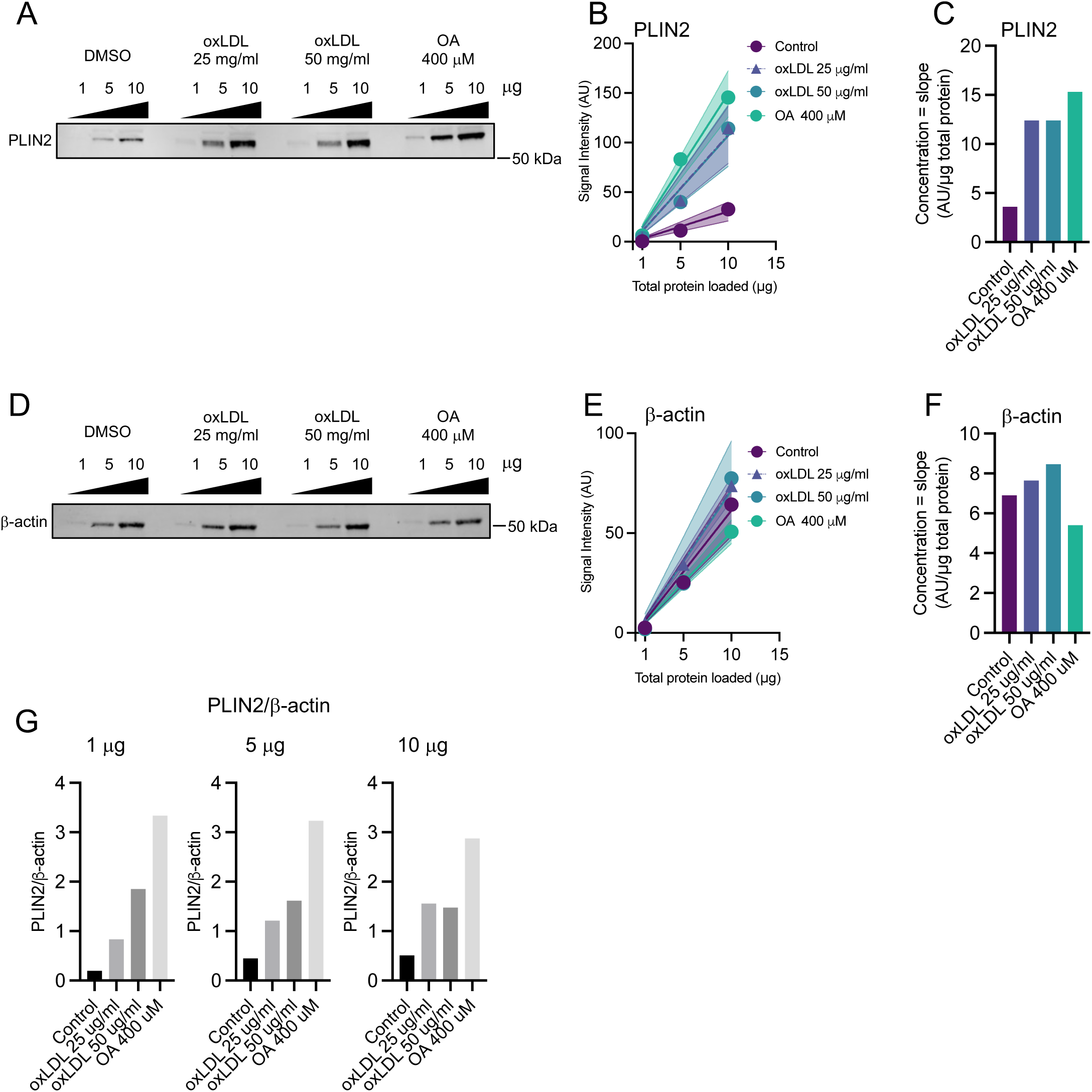
Accuracy of t-WB as compared to regular WB analysis. A) Representative blot of THP-1 cells treated with vehicle (DMSO), 25 μg/μl oxLDL, 50 μg/μl oxLDL or 400 μM OA. Each cell lysate sample was loaded at either 1, 5, 10 μg of total protein per well and PLIN2 protein was detected. B) t-WB linear regression plot for PLIN2. 95% CI are shown. C) PLIN2 concentration expressed as signal intensity per total μg protein loaded. D) Representative blot of β actin detected from the same samples and membranes as PLIN2 shown in A. E) t-WB linear regression plot for β actin. 95% CI are shown. F) β actin concentration expressed as signal intensity per total μg protein loaded. G) Classical WB analysis of PLIN2 expression using one single data point for quantification and the β actin signal to normalise between samples, shown separately for each of the loaded samples (1, 5 or 10 μg total loaded protein mass). PLIN2: perilipin 2; oxLDL: oxidized low-density lipoprotein; OA: bovine serum albumin (BSA)-coupled oleic acid; CI: confidence intervals.

To assess the precision of the t-WB method, we compared the signals from seven independent blots in which we determined the expression of low-density lipoprotein receptor (LDLR) in the same whole lysate specimen of HepG2 cells (Figure 3A-E, Supplementary Figure 1A-H). We used the t-WB to analyse the first derivatives obtained from the regression lines from the seven blots (Figure 3B-C; Supplementary Table 3A) and we calculated the coefficient of variation (CV) using the slopes, which was 33% (Figure 3C; Supplementary Table 3A). The CV, *i.e,* the standard deviation divided by the mean, was used instead of the standard deviation only, as it is insensitive to effects of variance in optical density, which can differ among different blots. To test whether the high variance among the signals could be reduced by using a single specific data point as internal control for the different blots, we randomly selected WB7 (Supplementary Table 3A) as a reference blot, and normalised all the signals in WB1 to WB6 by the signal detected at 20 μg, at 40 μg (Supplementary Figure 1A- D) or at 60 μg (Figure 3D-E) of total protein loaded in WB7 (Supplementary Table 3A). Thus, the respective 7 linear curves were plotted and LDLR concentration in AU/μg total protein loaded was re-estimated by the first derivatives. Performing this normalization reduced the inter-blot CV from 33% to 21.3% when the signal at 20 μg total protein loaded was used as internal control (Supplementary Figure 1A-B), to 11.7% when normalised for the signal at 40 μg (Supplementary Figure 1C-D), and to 4.5% when the signal at 60 μg was used (Figure 3D- E; Supplementary Table 3A). To confirm the reproducibility of this approach, we probed the same 7 membranes with a different primary antibody targeting the lysosome-associated membrane protein type 2A (LAMP2A) and performed the same analytical steps (Figure 3F-J; Supplementary Figure 1E-H, Supplementary Table 3B-C). Similarly, the inter-blot CV for the first derivatives of LAMP2A was reduced from 66% (Figure 3H) to 8.3%, when the signals in WB 1 to 6 were normalized for the signals of the 60 μg total protein loaded in WB7 (Figure 3J; Supplementary Table 3B). Finally, to show the robustness of the method and corroborate our results, we performed the same analysis using another lysate from HepG2 cells treated with atorvastatin (5 μM), which was analysed in 5 independent WBs and probed for LDLR (Supplementary Figure 1I-Q, Supplementary Table 3D) and LAMP2A (Supplementary Figure 1R-Z, Supplementary Table 3E-F). Once again, the normalization by the signal stemming from the highest concentration of loaded protein from one randomly chosen blot (WB1) reduced the CV from 13.2% (Supplementary Figure 1J-K) to 4.8% (Supplementary Figure 1P-Q) for LDLR, and from 67.2% (Supplementary Figure 1S-T) to 11.6% (Supplementary Figure 1Y-Z) for LAMP2A. We conclude that normalizing the signal intensity for an internal reference or internal control loaded at a single mass having a higher signal-to-background ratio, which in our experimental setting corresponds to 60 μg of total protein, allows a reliable comparison of results from multiple blots with a higher level of precision.

**Figure 3.**
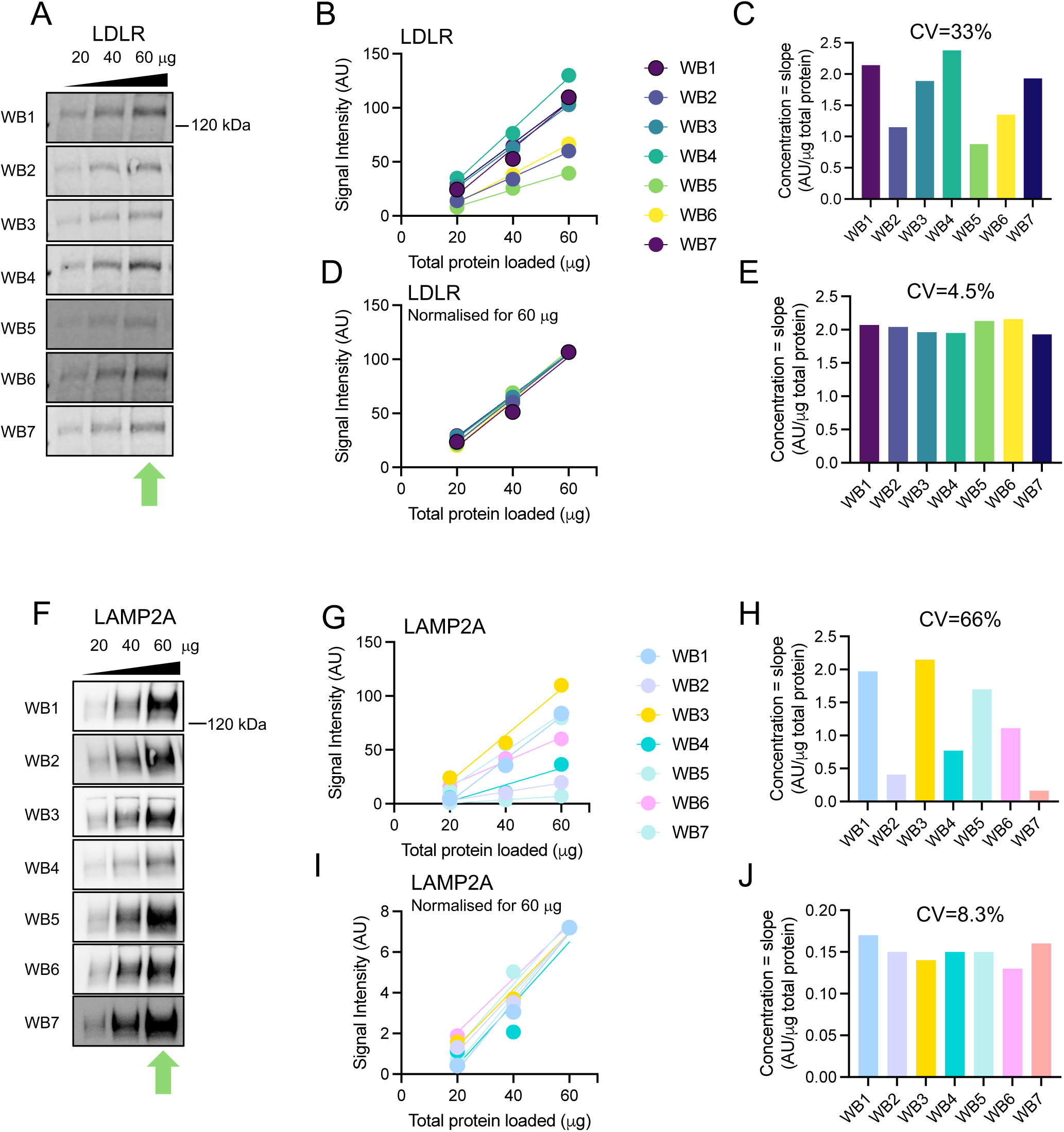
A single dilution reduces the coefficient of variations given by blot-to-blot variance. A) A single lysate of HepG2 cells treated with DMSO was loaded at 20, 40, 60 μg of total protein onto 7 different WB membranes and probed for the LDLR. B) t-WB regression plots for the LDLR signal on each of them. C) Calculated LDLR concentrations from each WB expressed as signal intensity per total μg loaded protein and CV of the concentrations. D) t- WB regression plots for LDLR after normalization of all data points using the WB7-60 μg data point (indicated in A by the green arrow). E) Calculated LDLR concentrations from each WB expressed as signal intensity per total μg loaded protein after normalization and CV of the resulting concentrations. F) Re-probing the same seven WB membranes shown in A) with LAMP2A antibody. G) t-WB regression plots for LAMP2A. H) Calculated LAMP2A concentrations from each WB expressed as signal intensity per total μg loaded protein and CV of the concentrations. I) t-WB regression plots for LAMP2A after normalization of all data points using the WB7-60 μg data point (green arrow in F). J) Calculated LAMP2A concentrations from each WB expressed as signal intensity per total μg loaded protein after normalization and CV of the resulting concentrations. LDLR: Low-density lipoprotein receptor; LAMP2A: lysosome-associated membrane protein type 2A; CV: coefficient of variation.

To further prove the robustness of t-WB, we challenged another commonly employed step in WB analysis: membranes are usually either cut and the fragments probed in parallel with different antibodies, or they are subjected to several runs of antibody probing, stripping, blocking and re-probing. To test whether t-WB could detect changes in protein concentrations upon such re-probing and could correct for it, we loaded the same titrated THP-1 cell lysate twice onto the same gel. After the standard run, transfer and blocking, the membrane was cut in two halves (P1 and P2), each bearing the same titrated sample (Figure 4A). P1 was probed directly for the cytosolic chaperone heat shock cognate 70 (HSC70; the main target protein for this experiment), whereas P2 was sequentially first probed for PLIN2 (not shown) and then for HSC70 (Figure 3A). When plotting the resulting protein signals, we could detect that the highest sample concentration (20 μg) on P1 was saturated, resulting in a R^2^ of 0.86 (Figure 4B, white point, Supplementary Table 4A). Thus we excluded this point from further analysis and continued with 3 points loading for P1 and 4 for P2 (Figure 4C). HSC70 protein concentration was thereafter assessed by t-WB (Figure 4D, Supplementary Table 4B). As expected, the HSC70 signal coming from the P2 membrane resulted in a signal with reduced intensity, due to the re-probing, which led to an underestimation of the concentration. As shown in Figure 4D, the HSC70 protein concentration calculated from P2 using the t-WB method was much lower than the one from P1 (13.5 *vs.* 4.9 AU/μg protein; Fig 4D, Supplementary Table 4B). We then applied the method of normalisation described above, using the signal of P2 at 10 μg as internal control and we reduced the CV from 65.9% (Figure 4D) to 3.7% (Figure 4F and Supplementary Table 4B). With this normalization, we obtained a comparable quantification of HSC70 protein concentration (13.5 AU/μg protein in P1 *vs.*12.8 AU/μg protein in P2), confirming the accuracy of t-WB and the benefits of using it in such experimental conditions.

**Figure 4.**
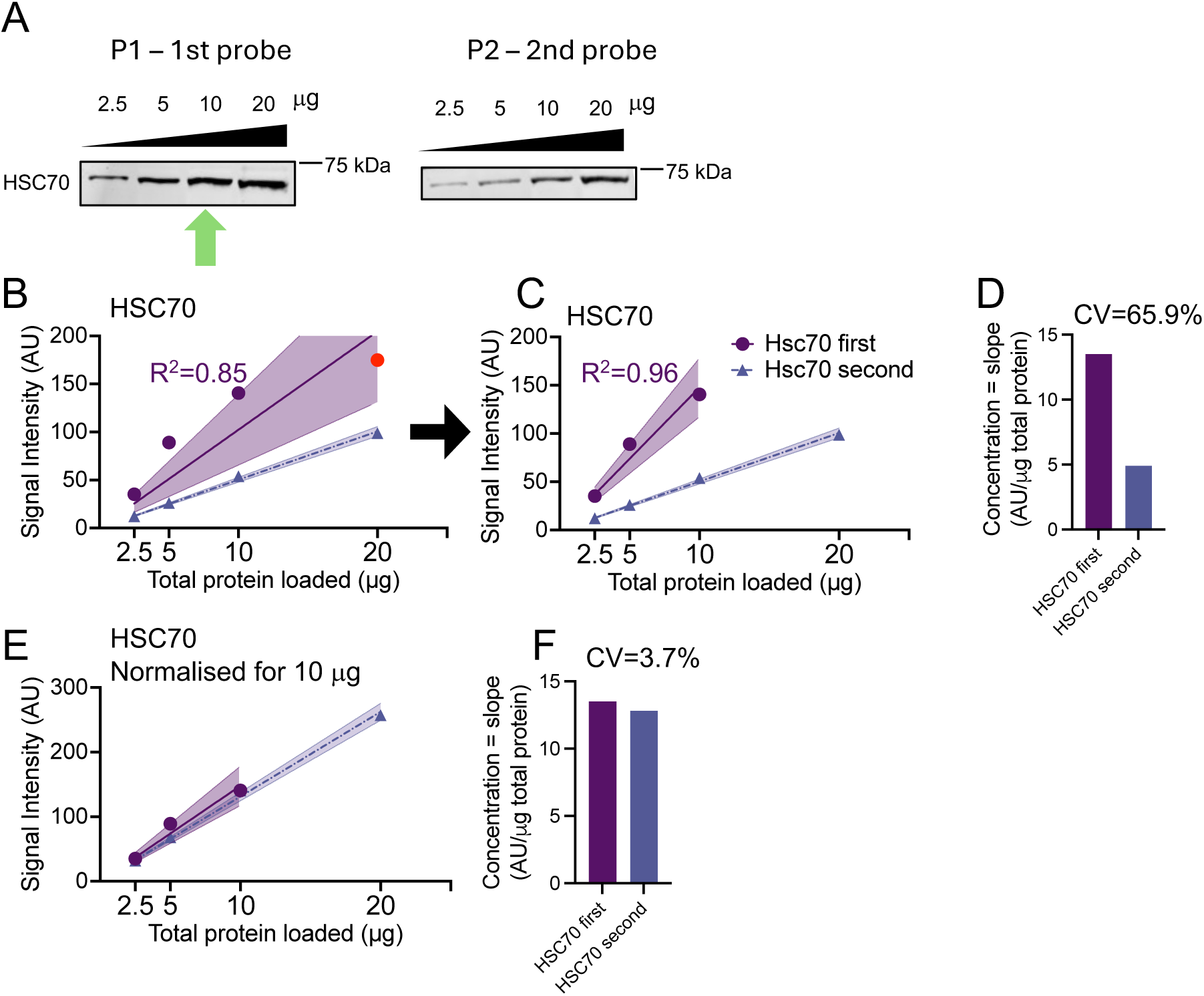
t-WB can be used to reduce the signal variability induced by re-probing. A) THP-1 cell lysate was loaded at 2.5, 5,10 and 20 μg of total protein mass in duplicate onto the same gel. After protein transfer and blocking the membrane was cut into two parts, P1 and P2: P1 was probed directly for HSC70, while P2 was first probed for PLIN2, stripped and subsequently probed for HSC70. B) t-WB regression plots of HSC70, purple dots show results from P1 (the red dot is the saturated P1 data point), while lilac dots show results from P2. 95% CI are shown. C) t-WB regression plots of HSC70 after exclusion of the saturated P1 data point. 95% CI are shown. D) Calculated concentration of HSC70 from each of the membranes and CV of the concentrations. E) t-WB regression plot after normalization of all points by the for P1-10 μg data point (indicated in A by a green arrow). 95% CI are shown. F) Calculated concentration of HSC70 from each WB expressed as signal intensity per total μg of loaded protein after normalization and CV of the resulting concentrations. HSC70: heat shock cognate 70; CI: confidence intervals; CV: coefficient of variation.

Finally, we wanted to prove if the use of t-WB with a single data point as internal control loaded at a single mass of total protein and independent from the sample tested, would allow the comparison of two samples run on two independent t-WBs (Figure 5). We loaded two gels (Gel 1 and Gel 2) with increasing amounts of total protein (1, 5, 10 μg) of cell lysates from THP-1 either incubated with OA (400 μM) or in basal conditions (control), together with an independent sample of THP-1 cell lysate loaded at 10 μg as internal control. After, gel electrophoresis and transfer onto membranes PLIN2 was immunodetected (Figure 5A). When plotting all signals in the same xy-graph, we observed that both signals and respective slope of the curves in Gel 1 were higher than those in Gel 2 (Figure 5B; Supplementary Table 5A-B). Despite this, we could show that in each independent WB, OA treatment led to a similar increase PLIN2 protein expression (AU/μg total protein) compared to control (4.9- and 4.5- fold increase in Gel 1 and Gel 2 respectively; Figure 5C, Supplementary Table 5B). Nevertheless, when we compared OA in Gel 2 *vs.* control in Gel 1 we observed a 21.9 fold increase in PLIN2 protein while no differences when between OA from Gel 1 *vs.* control from Gel 2 (Figure 5C, Supplementary Table 5B). We then normalised all signals detected in Gel 1 by the internal control, randomly choosing Gel 2 as reference (Supplementary Table 5B). As shown in Figure 5D the normalisation resulted in comparable linear functions respectively for both control and OA-treated samples within the 2 different WBs, and in turns allowed the inter WB comparison of the samples. As shown in Figure 5E and Supplementary Table 5B, we observed a 4.9-fold increase in PLIN2 protein comparing OA in Gel 2 *vs.* control in Gel 1 and a similar 4.7-fold increase when we compared OA from Gel 1 *vs.* control from Gel 2.

**Figure 5.**
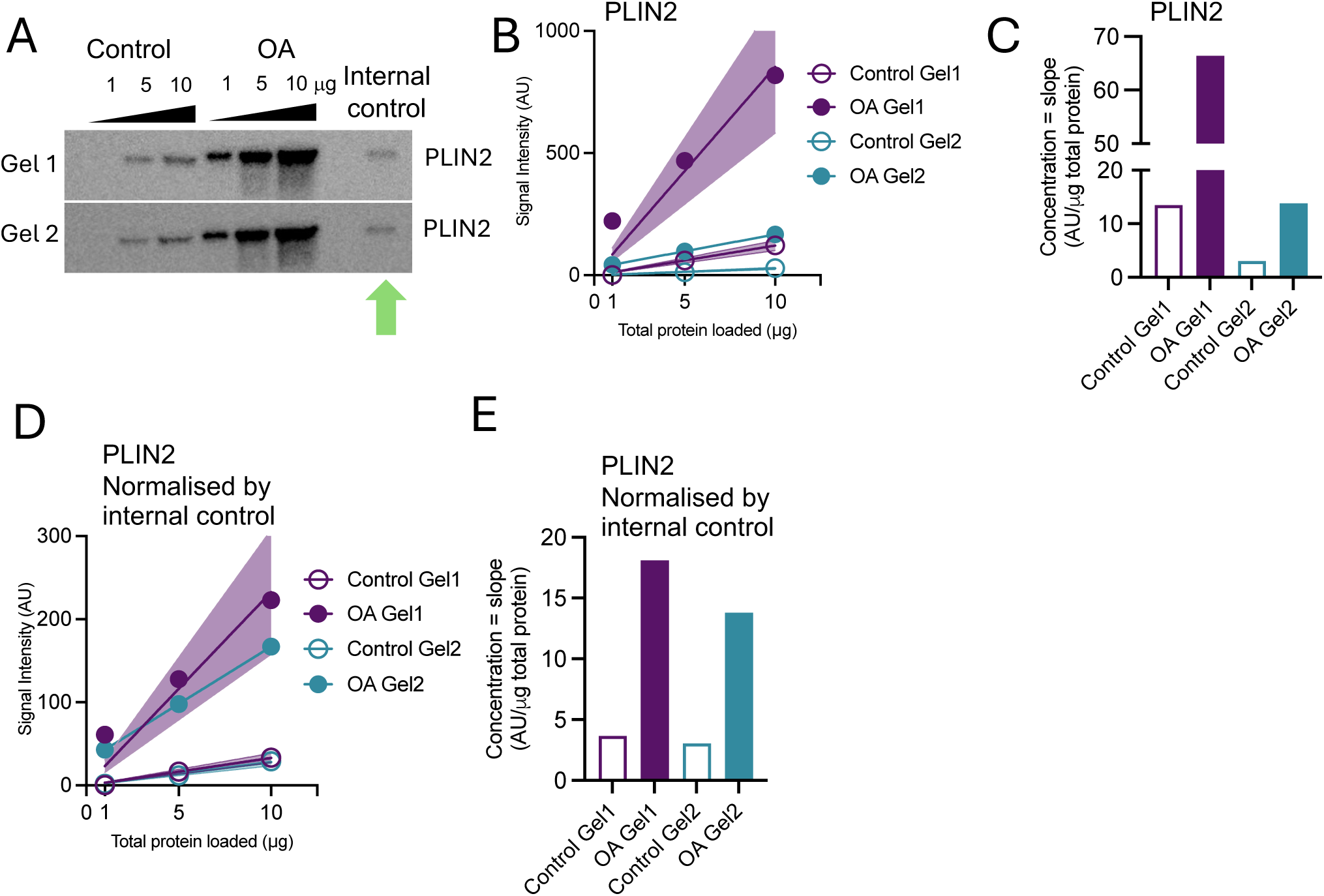
The internal control allows comparison of different samples from different t- WBs. A) THP-1 cell lysate control (left) and OA-treated (right) was loaded at 1, 5, 10 μg of total protein mass and a THP-1 internal control sample in two different gels: Gel 1 and Gel2. After protein transfer and blocking the membranes were probed for PLIN2; B) t-WB regression plots of PLIN2 from both Gel 1 and Gel 2. 95% CI are shown. C) Calculated concentrations of PLIN2 from Gel 1 and Gel 2 in control and OA treatment. D) t-WB regression plot of PLIN2 after normalization of all points by the internal control from Gel 2 (indicated in A by a green arrow). 95% CI are shown. E) Calculated concentration of PLIN2 from each WB expressed as signal intensity per total μg of loaded protein after normalization. PLIN2: perilipin 2; CI: confidence intervals; OA: bovine serum albumin (BSA)-coupled oleic acid.

## Discussion

The immunoblot, also known as Western blot, is one of the most widely utilized techniques in molecular biology for detecting and quantifying specific proteins within biological samples. While conceptually straightforward, its execution is complex and sensitive to various sources of bias. Consequently, immunoblots can be challenging to reproduce consistently and may yield inaccurate results if not rigorously validated through multiple repetitions. Here we have described the framework, the standardization, and the validation of the t-WB, allowing the accurate and quantitative determination of proteins of interest. Utilizing at least three sample concentrations, optimized within the linear range of signal detection, offers several advantages, *e.g.*: i) enables early detection of operator errors and other sources of bias that might be difficult to identify in a single point immunoblot experiment; ii) enhances the precision of protein quantification; and iii) significantly improves overall data accuracy.

Several researcher have suggested and recommended the use of calibration curves, dilutions and sample mixtures as a standard curves in Western blot experimental designs to improve the outcomes (Butler, Paul, Chan, Smith, & Tolosa, 2019; Charette, Lambert, Nadeau, & Landry, 2010; Pitre, Pan, Pruett, & Skalli, 2007; Suzuki et al., 2011; Taylor et al., 2013). However, the proposed approaches give relative semi-quantitative quantification and rely on housekeeping protein or whole protein signal normalization, which may lead to data misinterpretation has been extensively shown by our work and by others (McDonough, Veiras, Minas, & Ralph, 2015). Thus, our t-WB framework represents a significant advancement in terms of applicability, robustness, and precision by eliminating the reliance on housekeeping proteins for normalization.

The t-WB provides a range of precise, accurate and robust information that are neglected by classical WB techniques. First, the titration of the sample specimen and the consequent linear function with its R^2^ inform on operator errors and signal saturation for the protein of interest (Figure 1C-D). Moreover, the mathematical function of the linear regression bears another very useful advantage: the possibility to compare the intercept *b*. For an ideal blot (Figure 1B), all the *b-*values obtained either from the different samples run within in the same t-WB, or from the same sample run as different t-WB replicates, would be the same and approaching zero. This is assuming that: i) the background does not influence the signal detection; ii) the affinity/avidity of the antibody towards the protein of interest in the different specimens does not change within the same WB, or between replicate WBs; and iii) the loading of the specimen total protein mass is accurate. Pillai-Kastoori and colleagues described that the linear relationship between sample concentration and band intensity is lost at low and high concentration of protein loaded, since the tail and shoulder end of the intensity readout curve are strongly affected by noise and saturated signal, respectively (Pillai-Kastoori et al., 2020 (Pillai-Kastoori, Schutz-Geschwender, & Harford, 2020)). In line with this, Pitre and colleagues explained that the physical relation between the incident and emergent energy of light traversing a solid substance (*i.e.*, a WB band on a film) is logarithmic and not linear, due to the Beer-Lambert law (Pitre et al., 2007). Affinity and avidity are intrinsic properties of antibodies. Affinity refers to the strength of interaction between an antigen’s epitope and an antibody’s paratope at a single binding site, while avidity represents the sum of binding strengths across all binding sites. Theoretically, these properties should remain constant for a given antibody. However, in practice, the physical presence of co-localizing proteins on blot membranes can affect both affinity and avidity. This occurs through mechanisms such as steric hindrance, epitope masking, conformational changes, crowding effects, and non-specific interactions. This phenomenon is crucial to consider in protein analysis but is often overlooked. So, the varying b-values observed among different specimens (e.g., control *vs*. treated) analyzed in the same t-WB can serve as a valuable indicator of co-localizing proteins present in differing amounts. As we clearly showed (Figure 5), this effect does not affect the first derivative and thus can be disregarded when comparing samples within the same WB, but it appears to be critical when comparing protein expression between sample condition measure in different WB. This problem can be overcome in the t-WB by the use of an internal control loaded at one protein mass in all the several WB runs. The use of a single-point internal control to monitor and correct for signal variability, allowed indeed not only an accurate determination of protein concentration in same sample among different WBs, but also reduced the variation of results among repeated WBs.

To show the applicability of the t-WB to different laboratory setups and methodologies, we have performed the experiments using different WB techniques, from precast gels to homemade gels, wet transfer to semi-dry, nitrocellulose membranes and PVDF, chemiluminescence and fluorescent signal detection, and we detected signals using either Li-cor or ChemiDoc machines. In all different setups and settings, the application of the t-WB produced more accurate and less variable results than the classical WB.

Finally, a trade-off of t-WB is obviously the higher amount of biological material required to perform the titration, and the consequential higher number of blots per experiment needed. The payoff is however the generation of more solid and reproducible research, based on a more accurate, precise, and robust quantitative determination of protein expression, allowing also the ability to detect more subtle changes. For all these reasons we advocate for the systematic use of t-WB in science.

## Methods

### Cell culture and material

THP-1 were purchased from ATCC (American Type Culture Collection, Gaithersburg, MD, USA). RPMI 1640 medium with Glutamax supplement and heat inactivated foetal bovine serum (FBS) were purchased from Thermo Fisher Scientific (Europe BV, The Netherlands). Tissue culture flasks, plates, scrapers, and tubes were purchased from Thermo Fisher Scientific (Europe BV, The Netherlands) or Falcon (Lincoln, NY, USA). THP-1 cells were cultured in 6 well-plates, at 37°C in a 5% CO_2_ atmosphere in RPMI supplemented with 10% FBS and 50 ng/ml phorbol myristate acetate (PMA; Merck Life Science AB, Sweden) for 24h, then rested for 48h and treated with treatment media for additional 24h. Treatment media were culture media with 25-50 μg/ml of oxidized LDL or 400 μM of oleate. Cells were washed twice with cold PBS, then harvested using a scraper in lysis buffer (50 mM HEPES pH 8,0, 150 mM NaCl, 1% NP-40). Protein isolation with lysis buffer was performed for 15 min in ice, then centrifuges at 16,000 x g for 15 min at 4°C. The collected supernatant (total protein) was stored at -20°C. The total protein content was quantified by Pierce BCA Protein assay kit (Thermo Fisher Scientific Europe BV, The Netherlands).

Hepg2 were purchased from LGC Standards (Wesel, Germany). Dulbecco’s Modified Essential Medium (DMEM) with 1 g/L glucose and w/o pyruvate, Reduced-Serum Medium w/o phenol red (Opti-MEM), Trypsin-EDTA, Penicillin-Streptomycin (Pen-Strep) were purchased from Thermo Fisher Scientific (Europe BV, The Netherlands). Tissue culture flasks, plates, scrapers, and tubes were purchased from Thermo Fisher Scientific (Europe BV, The Netherlands), or Falcon (Lincoln, NY, USA). We previously developed a protocol that improves the phenotype of Hepg2 cells and make them to become more hepatocyte-like in term of hepatic lipid metabolism (hs-Hepg2; (Pramfalk, Larsson, Härdfeldt, Eriksson, & Parini, 2016). According to this protocol Hepg2 cells were cultured in 6 well-plates, at 37 °C in a 5% CO_2_ atmosphere in DMEM supplemented with 100 U/mL Pen + 100 μg/mL Strep and 2% human serum for 10 days. Then, hs-Hepg2 were incubated for 18h in Opti-MEM. Atorvastatin (5 μM) treatment was administered for 16h prior to harvest. Cells were washed in cold PBS, then harvested in RIPA buffer. Protein isolation with RIPA buffer was performed for 15 min in ice, then centrifuges at 16,000 x g for 15 min at 4°C. The collected supernatant (total protein) was stored at -80°C. The total protein content was quantified by Lowry method (DC^TM^ protein assay, Bio-Rad Laboratories, Hercules, USA).

### Western Blot for PLIN2 and HSC70

Protein lysates were diluted to a final concentration of 0.5 μg/ml and three total protein dilutions were prepared per sample per condition: for the PLIN2 WBs (Figure 2A and Figure 5A) 1, 5, 10 μg, for the HSC70 WB (Figure 4A) 2.5, 5, 10, 20 μg. Samples were loaded on 10% 18-well Stain free SDS-gel (Bio-Rad Laboratories, Hercules, USA) for electrophoresis, then transferred on low fluorescence PVDF membrane (Bio-Rad Laboratories, Hercules, USA) using the Turbo Transfer system. Membranes were blocked in 5% BSA in TBST, then probed for immunoblotting. For the PLIN2 WB, anti-PLIN2 antibody was diluted 1:1000 in 5% BSA in TBST (Origene). Secondaries Goat Anti-Rabbit IgG StarBright Blue 700 (Bio-Rad Laboratories, Hercules, USA) diluted 1:2500 for PLIN2 and Anti-Actin hFAB Rhodamine Antibody (Bio-Rad Laboratories, Hercules, USA) diluted 1:5000 in TBS were used. Signals are captured following antibody’s instructions using a ChemiDoc MP Imaging system with Image Lab Touch Software (Bio-Rad, Hercules, CA, USA).

For the HSC70 WB, anti-HSC70 antibody was diluted 1:1000 in 5% BSA in TBST (Novus Biologicals, Bio-Techne Ltd., UK). Secondary HRP-conjugated Goat Anti-Mouse IgG secondary antibody diluted 1:10000 in TBST (Bio-Rad Laboratories, Hercules, USA) were used. Blots were developed using enhanced chemiluminescence reagent kit (GE Healthcare Sverige AB, Sweden) and a ChemiDoc MP Imaging system with Image Lab Touch Software.

### Western Blot for LDLR and LAMP2A

Reduced pooled Hepg2 cell lysates (20, 40, 60 μg protein) were separated on a NuPage 3-8% Tris-Acetate gel and then transferred onto nitrocellulose membranes (Thermo Fisher Scientific Europe BV, The Netherlands). After blocking in StartingBlock™ T20 (TBS) Blocking Buffer (Thermo Fisher Scientific Europe BV, The Netherlands), the nitrocellulose membranes were incubated overnight at 4°C with polyclonal human LDL-Receptor Ab hosted in Rabbit (1:1000; LS-C146979; LS-Bio, Seattle, USA). After washing, membranes were incubated with IRDye^®^ 800CW Goat anti-Rabbit IgG Secondary Antibody (1:40000; 926- 32211, LI-COR Biosciences – GmbH, Germany). The specific bands for the LDL-Receptor glycosylated mature form (about 130 kDa) were detected by Odissey XF imaging system (LI-COR Biosciences – GmbH, Germany). The membranes were dried and kept in a plastic container at RT. The membranes were re-hydrated in PBS for 1h at RT before being incubated overnight at 4°C with polyclonal human LAMP2A antibody hosted in Rabbit (1:1000; ab18528; Abcam, USA). After washing, membranes were incubated with goat anti-rabbit IgG, (H + L) secondary antibody, HRP diluted 1:10000 in TBST (Thermo Fisher Scientific, 31460). Blots were developed using enhanced chemiluminescence reagent kit (GE Healthcare Sverige AB, Sweden) and a ChemiDoc MP Imaging system with Image Lab Touch Software.

## Authorship contribution statement

PP conceptualised the method. AM, OW and MO performed the experiments. AM and MP analysed the data. AM wrote the first version of the manuscript and PP, MP, EE and CEH edited, reviewed and finalise the final manuscript. EE and PP supervised the project.

## Acknowledgments

This work was supported by grants from Karolinska Institutet, Region Stockholm, Swedish Research Council, Swedish Heart and Lung Foundation.

## Data analysis and graphing

Graphs and data integration were prepared using GraphPad Prism (GraphPad Software Inc., CA, USA); t-WB pipeline was created with BioRender.com.

## References

1. Ahmed, O., Pramfalk, C., Pedrelli, M., Olin, M., Steffensen, K., Eriksson, M., & Parini, P. (2019). Genetic depletion of Soat2 diminishes hepatic steatosis via genes regulating de novo lipogenesis and by GLUT2 protein in female mice. Digestive and Liver Disease, 51(7), 1016–1022.

2. Bass, J. J., Wilkinson, D. J., Rankin, D., Phillips, B. E., Szewczyk, N. J., Smith, K., & Atherton, P. J. (2017). An overview of technical considerations for Western blotting applications to physiological research. Scand J Med Sci Sports, 27(1), 4–25. doi:10.1111/sms.12702

3. Butler, T. A. J., Paul, J. W., Chan, E. C., Smith, R., & Tolosa, J. M. (2019). Misleading Westerns: Common Quantification Mistakes in Western Blot Densitometry and Proposed Corrective Measures. Biomed Res Int, 2019, 5214821. doi:10.1155/2019/5214821

4. Charette, S. J., Lambert, H., Nadeau, P. J., & Landry, J. (2010). Protein quantification by chemiluminescent Western blotting: elimination of the antibody factor by dilution series and calibration curve. J Immunol Methods, 353(1-2), 148–150. doi:10.1016/j.jim.2009.12.007

5. Gnocchi, D., Ellis, E. C. S., Johansson, H., Eriksson, M., Bruscalupi, G., Steffensen, K. R., & Parini, P. (2020). Diiodothyronines regulate metabolic homeostasis in primary human hepatocytes by modulating mTORC1 and mTORC2 activity. Mol Cell Endocrinol, 499, 110604. doi:10.1016/j.mce.2019.110604

6. Goasdoue, K., Awabdy, D., Bjorkman, S. T., & Miller, S. (2016). Standard loading controls are not reliable for Western blot quantification across brain development or in pathological conditions. ELECTROPHORESIS, 37(4), 630–634. doi:10.1002/elps.201500385

7. Janes, K. A. (2015). An analysis of critical factors for quantitative immunoblotting. Sci Signal, 8(371), rs2. doi:10.1126/scisignal.2005966

8. Li, R., & Shen, Y. (2013). An old method facing a new challenge: re-visiting housekeeping proteins as internal reference control for neuroscience research. Life Sci, 92(13), 747–751. doi:10.1016/j.lfs.2013.02.014

9. McDonough, A. A., Veiras, L. C., Minas, J. N., & Ralph, D. L. (2015). Considerations when quantitating protein abundance by immunoblot. Am J Physiol Cell Physiol, 308(6), C426–433. doi:10.1152/ajpcell.00400.2014

10. McHenry, E. W., & Graham, M. L. (1935). Estimation of Ascorbic Acid by Titration. Nature, 135(3421), 871–872. doi:10.1038/135871b0

11. Minniti, M. E., Pedrelli, M., Vedin, L.-L., Delbès, A.-S., Denis, R. G. P., Öörni, K., . . . Parini, P. (2020). Insights From Liver-Humanized Mice on Cholesterol Lipoprotein Metabolism and LXR-Agonist Pharmacodynamics in Humans. Hepatology, 72(2), 656–670. doi:10.1002/hep.31052

12. Ntoukas, A., Niarchos, A., Tsika, A. C., Mantzoukas, S., Spyroulias, G. A., & Poulas, K. (2021). A quantitative western blot technique using TMB: Comparison with the conventional technique. ELECTROPHORESIS, 42(6), 786–792. doi:10.1002/elps.202000306

13. Parini, P., Gustafsson, U., Davis, M. A., Larsson, L., Einarsson, C., Wilson, M., Sahlin, S. (2008). Cholesterol synthesis inhibition elicits an integrated molecular response in human livers including decreased ACAT2. Arteriosclerosis, thrombosis, and vascular biology, 28(6), 1200–1206.

14. Pedrelli, M., Davoodpour, P., Degirolamo, C., Gomaraschi, M., Graham, M., Ossoli, A., . . . Steffensen, K. R. (2014). Hepatic ACAT2 knock down increases ABCA1 and modifies HDL metabolism in mice. PLoS One, 9(4), e93552.

15. Pillai-Kastoori, L., Schutz-Geschwender, A. R., & Harford, J. A. (2020). A systematic approach to quantitative Western blot analysis. Anal Biochem, 593, 113608. doi:10.1016/j.ab.2020.113608

16. Pitre, A., Pan, Y., Pruett, S., & Skalli, O. (2007). On the use of ratio standard curves to accurately quantitate relative changes in protein levels by Western blot. Anal Biochem, 361(2), 305–307. doi:10.1016/j.ab.2006.11.008

17. Pramfalk, C., Ahmed, O., Pedrelli, M., Minniti, M. E., Luquet, S., Denis, R. G., Steffensen, K. R. (2022). Soat2 ties cholesterol metabolism to β-oxidation and glucose tolerance in male mice. Journal of Internal Medicine, 292(2), 296–307.

18. Pramfalk, C., Jakobsson, T., Verzijl, C. R., Minniti, M. E., Obensa, C., Ripamonti, F., Parini, P. (2020). Generation of new hepatocyte-like in vitro models better resembling human lipid metabolism. Biochimica et Biophysica Acta (BBA)-Molecular and Cell Biology of Lipids, 1865(6), 158659.

19. Pramfalk, C., Larsson, L., Härdfeldt, J., Eriksson, M., & Parini, P. (2016). Culturing of HepG2 cells with human serum improve their functionality and suitability in studies of lipid metabolism. Biochimica et Biophysica Acta (BBA) - Molecular and Cell Biology of Lipids, 1861(1), 51–59. doi:10.1016/j.bbalip.2015.10.008

20. Suzuki, O., Koura, M., Noguchi, Y., Uchio-Yamada, K., & Matsuda, J. (2011). Use of sample mixtures for standard curve creation in quantitative western blots. Experimental Animals, 60(2), 193–196.

21. Taylor, S. C., Berkelman, T., Yadav, G., & Hammond, M. (2013). A Defined Methodology for Reliable Quantification of Western Blot Data. Molecular Biotechnology, 55(3), 217–226. doi:10.1007/s12033-013-9672-6

22. Towbin, H., Staehelin, T., & Gordon, J. (1979). Electrophoretic transfer of proteins from polyacrylamide gels to nitrocellulose sheets: procedure and some applications. Proceedings of the National Academy of Sciences, 76(9), 4350–4354. doi:10.1073/pnas.76.9.4350

