## Supplementary Table 1-5 for "Titration-WB: A methodology for accurate quantitative protein determination overcoming reproducibility errors"

## A

| Figure 1B WB | |
|---|---|
| Protein loaded (µg) | Signal intensity (AU) |
| 2 | 10 |
| 8 | 40 |
| 20 | 100 |

## B

| Figure 1C WB | |
|---|---|
| Protein loaded (µg) | Signal intensity (AU) |
| 2 | 10 |
| 8 | 8 |
| 20 | 100 |

## C

| Figure 1D WB | |
|---|---|
| Protein loaded (µg) | Signal intensity (AU) |
| 2 | 25 |
| 8 | 118 |
| 20 | 120 |

## D

| Figure 1E WB | | |
|---|---|---|
| | Reduced avidity | Normal |
| Protein loaded (µg) | Signal intensity (AU) | Signal intensity (AU) |
| 0 | -1,167 | 2,833 |
| 2 | 2 | 6 |
| 4 | 4,5 | 8,5 |
| 6 | 8 | 12 |

## E

| Figure 1F WB | | | |
|---|---|---|---|
| | Correct loading | Corrected overstimation | Overstimation |
| Protein loaded (µg) | Signal intensity (AU) | Signal intensity (AU) | Signal intensity (AU) |
| 0 | -1,167 | 2,833 | 1,333 |
| 2 | 2 | 6 | |
| 4 | 4,5 | 8,5 | |
| 6 | 8 | 12 | |
| 3 | | | 6 |
| 5 | | | 8,5 |
| 7 | | | 12 |

### Supplementary Table 2

## A

| PLIN2 | Signal (AU) | | | Equation | Slope | Fold change |
|---|---|---|---|---|---|---|
| | Protein loaded (μg) | | | | | |
| | 1 | 5 | 10 | | | |
| Control | 513086 | 11346718 | 32711544 | y = 3 606 104,75x - 4 375 442,69 | 3606104,75 | 1,0 |
| oxLDL 25 μg/ml | 4143280 | 41508566 | 114533016 | y = 12 361 401,79x - 12 532 522,20 | 12375026,3 | 3,4 |
| oxLDL 50 μg/ml | 3853706 | 40005164 | 114275468 | y = 12 375 026,31x - 13 288 694,33 | 12361401,8 | 3,4 |
| OA 400 μM | 6439914 | 83306862 | 145565004 | y = 15 335 117,31x - 3 350 032,33 | 15335117,3 | 4,3 |

## B

| PLIN2 | Signal (AU)/10^6 | | | Equation | Slope | Fold change |
|---|---|---|---|---|---|---|
| | Protein loaded (μg) | | | | | |
| | 1 | 5 | 10 | | | |
| Control | 0,5 | 11,3 | 32,7 | y = 3,6061x - 4,3754 | 3,6 | 1,0 |
| oxLDL 25 μg/ml | 4,1 | 41,5 | 114,5 | y = 12,361x - 12,533 | 12,4 | 3,4 |
| oxLDL 50 μg/ml | 3,9 | 40,0 | 114,3 | y = 12,375x - 13,289 | 12,4 | 3,4 |
| OA 400 μM | 6,4 | 83,3 | 145,6 | y = 15,335x - 3,35 | 15,3 | 4,3 |

## C

| β-actin | Signal (AU) | | | Equation | Slope | Fold change |
|---|---|---|---|---|---|---|
| | Protein loaded (μg) | | | | | |
| | 1 | 5 | 10 | | | |
| Control | 2584860 | 25395412 | 64332596 | y = 6 898 834,03x - 6 022 825,51 | 6898834 | 1,0 |
| oxLDL 25 μg/ml | 4968050 | 34226786 | 73633402 | y = 7 639 804,85x - 3 136 213,21 | 7639804 | 1,1 |
| oxLDL 50 μg/ml | 2079278 | 24794422 | 77421448 | y = 8 459 633,08x - 10 352 993,77 | 8459633 | 1,2 |
| OA 400 μM | 1929486 | 25794196 | 50661112 | y = 5 396 541,43x - 2 653 289,61 | 5396541 | 0,8 |

## D

| β-actin | Signal (AU)/10^6 | | | Equation | Slope | Fold change |
|---|---|---|---|---|---|---|
| | Protein loaded (μg) | | | | | |
| | 1 | 5 | 10 | | | |
| Control | 2,6 | 25,4 | 64,3 | y = 6,8988x - 6,0228 | 6,90 | 1,0 |
| oxLDL 25 μg/ml | 5,0 | 34,2 | 73,6 | y = 7,6398x - 3,1362 | 7,64 | 1,1 |
| oxLDL 50 μg/ml | 2,1 | 24,8 | 77,4 | y = 8,4596x - 10,353 | 8,46 | 1,2 |
| OA 400 μM | 1,9 | 25,8 | 50,7 | y = 5,3965x - 2,6533 | 5,40 | 0,8 |

## E

| PLIN2 | PLIN2/β-actin | | | Fold change 1μg | Fold change 5μg | Fold change 10μg |
|---|---|---|---|---|---|---|
| | Protein loaded (μg) | | | | | |
| | 1 | 5 | 10 | | | |
| Control | 0,20 | 0,45 | 0,51 | 1,00 | 1,00 | 1,00 |
| oxLDL 25 μg/ml | 0,83 | 1,21 | 1,56 | 4,20 | 2,71 | 3,06 |
| oxLDL 50 μg/ml | 1,85 | 1,61 | 1,48 | 9,34 | 3,61 | 2,90 |
| OA 400 μM | 3,34 | 3,23 | 2,87 | 16,81 | 7,23 | 5,65 |

### Supplementary Table 3

## A

| WB # | Signal (AU) | | | Equation | Slopes | Ratio intergel 20µg | Signal adjusted at 20µg | | | Slopes | Ratio intergel 40µg | Signal adjusted at 40µg | | | Slopes | Ratio intergel 60µg | Signal adjusted at 60µg | | | Equation | Slopes |
|---|---|---|---|---|---|---|---|---|---|---|---|---|---|---|---|---|---|---|---|---|---|
| LDLR control | | | | | | | | | | | | | | | | | | | | | |
| | Protein loaded (µg) | | | | | | Protein loaded (µg) | | | | | Protein loaded (µg) | | | | | Protein loaded (µg) | | | | |
| | 20 | 40 | 60 | | | | 20 | 40 | 60 | | | 20 | 40 | 60 | | | 20 | 40 | 60 | | |
| 1 | 24,34 | 52,78 | 109,75 | y=2,1353x-23,122 | 2,14 | 1,21 | 29,48 | 63,94 | 132,94 | 2,59 | 1,22 | 29,78 | 64,59 | 134,30 | 2,61 | 0,97 | 23,64 | 51,27 | 106,60 | y = 2,074x - 22,458 | 2,07 |
| 2 | 14,00 | 33,92 | 60,01 | y=1,1501x-10,03 | 1,15 | 2,11 | 29,48 | 71,43 | 126,35 | 2,42 | 1,90 | 26,66 | 64,59 | 114,26 | 2,19 | 1,78 | 24,87 | 60,26 | 106,60 | y = 2,0432x - 17,818 | 2,04 |
| 3 | 27,27 | 62,63 | 102,75 | y=1,887x-11,263 | 1,89 | 1,08 | 29,48 | 67,70 | 111,07 | 2,04 | 1,03 | 28,13 | 64,59 | 105,98 | 1,95 | 1,04 | 28,29 | 64,97 | 106,60 | y = 1,9576x - 11,685 | 1,96 |
| 4 | 34,85 | 76,42 | 129,98 | y=2,378x-14,706 | 2,38 | 0,85 | 29,48 | 64,63 | 109,93 | 1,93 | 0,85 | 29,46 | 64,59 | 109,87 | 2,01 | 0,82 | 28,58 | 62,67 | 106,60 | y = 1,9503x - 12,061 | 1,95 |
| 5 | 7,95 | 25,51 | 39,38 | y=0,7857x-7,1457 | 0,88 | 3,71 | 29,48 | 94,52 | 145,94 | 2,91 | 2,53 | 20,15 | 64,59 | 99,73 | 1,99 | 2,71 | 21,53 | 69,04 | 106,60 | y = 2,1266x - 19,341 | 2,13 |
| 6 | 12,58 | 38,03 | 66,67 | y=1,3522x-14,992 | 1,35 | 2,34 | 29,48 | 89,12 | 156,20 | 3,17 | 1,70 | 21,37 | 64,59 | 113,22 | 2,30 | 1,60 | 20,12 | 60,81 | 106,60 | y = 2,162x - 23,97 | 2,16 |
| 7 | 29,48 | 64,59 | 106,60 | y=1,9279x-10,228 | 1,93 | 1,00 | 29,48 | 64,59 | 106,60 | 1,93 | 1,00 | 29,48 | 64,59 | 106,60 | 1,93 | 1,00 | 29,48 | 64,59 | 106,60 | y = 1,9279x - 10,228 | 1,93 |

| | Coefficient of variation |
|---|---|
| Slopes not normalised | 33,03 |
| Slopes adjusted at 20µg | 20,31 |
| Slopes adjusted at 40µg | 11,65 |
| Slopes adjusted at 60µg | 4,53 |

## B

| WB # | Signal (AU) | | | Equation | Slopes | Ratio intergel 20µg | Signal adjusted at 20µg | | | Slopes | Ratio intergel 40µg | Signal adjusted at 40µg | | | Slopes | Ratio intergel 60µg | Signal adjusted at 60µg | | | Slopes |
|---|---|---|---|---|---|---|---|---|---|---|---|---|---|---|---|---|---|---|---|---|
| LAMP2A control | | | | | | | | | | | | | | | | | | | | |
| | Protein loaded (µg) | | | | | | Protein loaded (µg) | | | | | Protein loaded (µg) | | | | | Protein loaded (µg) | | | |
| | 20 | 40 | 60 | | | | 20 | 40 | 60 | | | 20 | 40 | 60 | | | 20 | 40 | 60 | |
| 1 | 4870170 | 35635149 | 83715548 | y=1971134,45x-37438422,33 | 1971134 | 0,15 | 709410 | 5190770 | 12194368 | 287124 | 0,10 | 469975 | 3438820 | 8078616 | 190216 | 0,09 | 418695 | 3063602 | 7197138 | 169461 |
| 2 | 3625168 | 9757440 | 19875874 | y=406267,65x-5164545,33 | 406268 | 0,20 | 709410 | 1909436 | 3889515 | 79503 | 0,35 | 1277620 | 3438820 | 7004865 | 143181 | 0,36 | 1312689 | 3533210 | 7197138 | 147111 |
| 3 | 24166920 | 56591988 | 109975294 | y=2145209,35x-22230306,67 | 2145209 | 0,03 | 709410 | 1661235 | 3228280 | 62972 | 0,06 | 1468506 | 3438820 | 6682664 | 130354 | 0,07 | 1581561 | 3703562 | 7197138 | 140389 |
| 4 | 5716648 | 10495160 | 36472191 | y=768888,58x-13194210 | 768889 | 0,12 | 709410 | 1302402 | 4526033 | 95416 | 0,33 | 1873104 | 3438820 | 11950394 | 251932 | 0,20 | 1128079 | 2071033 | 7197138 | 151726 |
| 5 | 11509035 | 55712823 | 79694496 | y=1704636,53x19213345 | 1704637 | 0,06 | 709410 | 3434105 | 4912321 | 105073 | 0,06 | 710384 | 3438820 | 4919066 | 105217 | 0,09 | 1039371 | 5031375 | 7197138 | 153944 |
| 6 | 15822862 | 41962698 | 60131539 | y=1107716,93x5002977,33 | 1107717 | 0,04 | 709410 | 1881376 | 2695967 | 49664 | 0,08 | 1296675 | 3438820 | 4927747 | 90777 | 0,12 | 1893837 | 5022511 | 7197138 | 132583 |
| 7 | 709410 | 3438820 | 7197138 | y=162193,2x-2705938,67 | 162193 | 1,00 | 709410 | 3438820 | 7197138 | 162193 | 1,00 | 709410 | 3438820 | 7197138 | 162193 | 1,00 | 709410 | 3438820 | 7197138 | 162193 |

| | Coefficient of variation |
|---|---|
| Slopes not normalised | 65,97 |
| Slopes adjusted at 20µg | 68,20 |
| Slopes adjusted at 40µg | 35,72 |
| Slopes adjusted at 60µg | 8,30 |

## C

| WB # | Signal (AU)/10^6 | | | Equation | Slopes | Ratio intergel 20µg | Signal adjusted at 20µg | | | Slopes | Ratio intergel 40µg | Signal adjusted at 40µg | | | Slopes | Ratio intergel 60µg | Signal adjusted at 60µg | | | Slopes |
|---|---|---|---|---|---|---|---|---|---|---|---|---|---|---|---|---|---|---|---|---|
| LAMP2A control | | | | | | | | | | | | | | | | | | | | |
| | Protein loaded (µg) | | | | | | Protein loaded (µg) | | | | | Protein loaded (µg) | | | | | Protein loaded (µg) | | | |
| | 20 | 40 | 60 | | | | 20 | 40 | 60 | | | 20 | 40 | 60 | | | 20 | 40 | 60 | |
| 1 | 4,87 | 35,64 | 83,72 | y=1,97x-37,44 | 1,97 | 0,15 | 0,71 | 5,19 | 12,19 | 0,29 | 0,10 | 0,47 | 3,44 | 8,08 | 0,19 | 0,09 | 0,42 | 3,06 | 7,20 | 0,17 |
| 2 | 3,63 | 9,76 | 19,88 | y=0,41x-5,16 | 0,41 | 0,20 | 0,71 | 1,91 | 3,89 | 0,08 | 0,35 | 1,28 | 3,44 | 7,00 | 0,14 | 0,36 | 1,31 | 3,53 | 7,20 | 0,15 |
| 3 | 24,17 | 56,59 | 109,98 | y=2,15x-22,23 | 2,15 | 0,03 | 0,71 | 1,66 | 3,23 | 0,06 | 0,06 | 1,47 | 3,44 | 6,68 | 0,13 | 0,07 | 1,58 | 3,70 | 7,20 | 0,14 |
| 4 | 5,72 | 10,50 | 36,47 | y=0,77x-13,19 | 0,77 | 0,12 | 0,71 | 1,30 | 4,53 | 0,10 | 0,33 | 1,87 | 3,44 | 11,95 | 0,25 | 0,20 | 1,13 | 2,07 | 7,20 | 0,15 |
| 5 | 11,51 | 55,71 | 79,69 | y=1,70x19,21 | 1,70 | 0,06 | 0,71 | 3,43 | 4,91 | 0,11 | 0,06 | 0,71 | 3,44 | 4,92 | 0,11 | 0,09 | 1,04 | 5,03 | 7,20 | 0,15 |
| 6 | 15,82 | 41,96 | 60,13 | y=1,11x-5 | 1,11 | 0,04 | 0,71 | 1,88 | 2,70 | 0,05 | 0,08 | 1,30 | 3,44 | 4,93 | 0,09 | 0,12 | 1,89 | 5,02 | 7,20 | 0,13 |
| 7 | 0,71 | 3,44 | 7,20 | y=0,16x-2,70 | 0,16 | 1,00 | 0,71 | 3,44 | 7,20 | 0,16 | 1,00 | 0,71 | 3,44 | 7,20 | 0,16 | 1,00 | 0,71 | 3,44 | 7,20 | 0,16 |

## D

| WB # | Signal (AU) | | | Equation | Slopes | Ratio intergel 20µg | Signal adjusted at 20µg | | | Slopes | Ratio intergel 40µg | Signal adjusted at 40µg | | | Slopes | Ratio intergel 60µg | Signal adjusted at 60µg | | | Slopes |
|---|---|---|---|---|---|---|---|---|---|---|---|---|---|---|---|---|---|---|---|---|
| LDLR atorva | | | | | | | | | | | | | | | | | | | | |
| | Protein loaded (µg) | | | | | | Protein loaded (µg) | | | | | Protein loaded (µg) | | | | | Protein loaded (µg) | | | |
| | 20 | 40 | 60 | | | | 20 | 40 | 60 | | | 20 | 40 | 60 | | | 20 | 40 | 60 | |
| 1 | 70,16 | 224,73 | 350,56 | y = 7,01x - 65,249 | 7,01 | 1,00 | 70,16 | 224,73 | 350,56 | 7,01 | 1,00 | 70 | 225 | 351 | 7,01 | 1,00 | 70 | 225 | 351 | 7,01 |
| 2 | 70,01 | 192,88 | 318,47 | y = 6,2117x - 54,684 | 6,2117 | 1,00 | 70,16 | 193,30 | 319,17 | 6,2253 | 1,17 | 82 | 225 | 371 | 7,2377 | 1,10 | 77 | 212 | 351 | 6,8375 |
| 3 | 53,41 | 179,90 | 271,66 | y = 5,4562x - 49,924 | 5,4562 | 1,31 | 70,16 | 236,31 | 356,83 | 7,1669 | 1,25 | 67 | 225 | 339 | 6,8159 | 1,29 | 69 | 232 | 351 | 7,0408 |
| 4 | 76,69 | 186,57 | 285,33 | y = 5,216x - 25,774 | 5,216 | 0,91 | 70,16 | 170,68 | 261,02 | 4,7716 | 1,20 | 92 | 225 | 344 | 6,2828 | 1,23 | 94 | 229 | 351 | 6,4084 |
| 5 | 78,86 | 186,19 | 289,81 | y = 5,2736x - 25,991 | 5,2736 | 0,89 | 70,16 | 165,64 | 257,82 | 4,6915 | 1,21 | 95 | 225 | 350 | 6,3654 | 1,21 | 95 | 225 | 351 | 6,3791 |

| | Coefficient of variation |
|---|---|
| Slopes not normalised | 13,18 |
| Slopes adjusted at 20µg | 19,90 |
| Slopes adjusted at 40µg | 6,10 |
| Slopes adjusted at 60µg | 4,77 |

## E

| WB # | Signal (AU) | | | Equation | Slopes | Ratio intergel 20µg | Signal adjusted at 20µg | | | Slopes | Ratio intergel 40µg | Signal adjusted at 40µg | | | Slopes | Ratio intergel 60µg | Signal adjusted at 60µg | | | Slopes |
|---|---|---|---|---|---|---|---|---|---|---|---|---|---|---|---|---|---|---|---|---|
| LAMP2A atorva | | | | | | | | | | | | | | | | | | | | |
| | Protein loaded (µg) | | | | | | Protein loaded (µg) | | | | | Protein loaded (µg) | | | | | Protein loaded (µg) | | | |
| | 20 | 40 | 60 | | | | 20 | 40 | 60 | | | 20 | 40 | 60 | | | 20 | 40 | 60 | |
| 1 | 528360 | 3797938 | 6406394 | y=146950,85-2300470 | 146950,85 | 1,00 | 528360 | 3797938 | 6406394 | 146950,85 | 1,00 | 528360 | 3797938 | 6406394 | 146950,85 | 1,00 | 528360 | 3797938 | 6406394 | 146950,85 |
| 2 | 11658680 | 26568768 | 49688268 | y=950739,78724349,33 | 950739,7 | 0,05 | 528360 | 1204071 | 2251824 | 43086,6 | 0,14 | 1666579 | 3797938 | 7102812 | 135905,83 | 0,13 | 1503174 | 3425557 | 6406394 | 122580,51 |
| 3 | 20771799 | 52867785 | 94473416 | y=1842540,43-17663950,33 | 1842540,43 | 0,03 | 528360 | 1344767 | 2403065 | 46867,61 | 0,07 | 1492213 | 3797938 | 6786821 | 132365,19 | 0,07 | 1408569 | 3585049 | 6406394 | 124945,62 |
| 4 | 17094942 | 34168420 | 53926530 | y=920789,7x1768290,67 | 920789,7 | 0,03 | 528360 | 1056057 | 1666728 | 28459,2 | 0,11 | 1900162 | 3797938 | 5994120 | 102348,96 | 0,12 | 2030854 | 4059159 | 6406394 | 109388,49 |
| 5 | 10970869 | 24884241 | 38702914 | y=693301,13-2879370,33 | 693301,13 | 0,05 | 528360 | 1198432 | 1863943 | 33389,57 | 0,15 | 1674420 | 3797938 | 5907002 | 105814,55 | 0,17 | 1815980 | 4119025 | 6406394 | 114760,36 |

| | Coefficient of variation |
|---|---|
| Slopes not normalised | 67,24 |
| Slopes adjusted at 20µg | 82,51 |
| Slopes adjusted at 40µg | 15,72 |
| Slopes adjusted at 60µg | 11,63 |

## F

| WB # | Signal (AU)/10^6 | | | Equation | Slopes | Ratio intergel 20µg | Signal adjusted at 20µg | | | Slopes | Ratio intergel 40µg | Signal adjusted at 40µg | | | Slopes | Ratio intergel 60µg | Signal adjusted at 60µg | | | Slopes |
|---|---|---|---|---|---|---|---|---|---|---|---|---|---|---|---|---|---|---|---|---|
| LAMP2A atorva | | | | | | | | | | | | | | | | | | | | |
| | Protein loaded (µg) | | | | | | Protein loaded (µg) | | | | | Protein loaded (µg) | | | | | Protein loaded (µg) | | | |
| | 20 | 40 | 60 | | | | 20 | 40 | 60 | | | 20 | 40 | 60 | | | 20 | 40 | 60 | |
| 1 | 0,53 | 3,80 | 6,41 | y = 0,147x - 2,3005 | 0,147 | 1,00 | 0,53 | 3,80 | 6,41 | 0,15 | 1,00 | 0,53 | 3,80 | 6,41 | 0,15 | 1,00 | 0,53 | 3,80 | 6,41 | 0,15 |
| 2 | 11,66 | 26,57 | 49,69 | y = 0,9507x - 8,7243 | 0,95 | 0,05 | 0,53 | 1,20 | 2,25 | 0,04 | 0,14 | 1,67 | 3,80 | 7,10 | 0,14 | 0,13 | 1,50 | 3,43 | 6,41 | 0,12 |
| 3 | 20,77 | 52,87 | 94,47 | y = 1,8425x - 17,664 | 1,84 | 0,03 | 0,53 | 1,34 | 2,40 | 0,05 | 0,07 | 1,49 | 3,80 | 6,79 | 0,13 | 0,07 | 1,41 | 3,59 | 6,41 | 0,12 |
| 4 | 17,09 | 34,17 | 53,93 | y = 0,9208x - 1,7683 | 0,92 | 0,03 | 0,53 | 1,06 | 1,67 | 0,03 | 0,11 | 1,90 | 3,80 | 5,99 | 0,10 | 0,12 | 2,03 | 4,06 | 6,41 | 0,11 |
| 5 | 10,97 | 24,88 | 38,70 | y = 0,6933x - 2,8794 | 0,69 | 0,05 | 0,53 | 1,20 | 1,86 | 0,03 | 0,15 | 1,67 | 3,80 | 5,91 | 0,11 | 0,17 | 1,82 | 4,12 | 6,41 | 0,11 |

### Supplementary Table 4

## A

| WB # | Signal (AU) | | | | Equation | R^2 |
|---|---|---|---|---|---|---|
| | Protein loaded (µg) | | | | | |
| | 2,5 | 5 | 10 | 20 | | |
| P1 | 35,20 | 89,13 | 140,64 | 174,94 | y = 7,3166x + 41,383 | 0,86 |
| P2 | 12,27 | 26,16 | 53,96 | 98,69 | y = 4,9171x + 1,6744 | 1,00 |

## B

| WB # | Signal (AU) | | | | Equation | Slope | Ratio intergel 2,5mg | Signal adjusted at 2,5µg | | | | Slope | Ratio intergel 5ug | Signal adjusted at 5µg | | | | Slope | Ratio intergel 10ug | Signal adjusted at 10µg | | | | Slope |
|---|---|---|---|---|---|---|---|---|---|---|---|---|---|---|---|---|---|---|---|---|---|---|---|---|
| | Protein loaded (µg) | | | | | | | Protein loaded (µg) | | | | | | Protein loaded (µg) | | | | | | Protein loaded (µg) | | | | |
| | 2,5 | 5 | 10 | 20 | | | | 2,5 | 5 | 10 | 20 | | | 2,5 | 5 | 10 | 20 | | | 2,5 | 5 | 10 | 20 | |
| P1 | 35,20 | 89,13 | 140,64 | | y = 13,522x + 9,4455 | 13,50 | 0,35 | 12,27 | 31,06 | 49,01 | | 13,5 | 0,29 | 10,33 | 26,16 | 41,28 | | 13,5 | 0,38 | 13,51 | 34,20 | 53,96 | | 13,5 |
| P2 | 12,27 | 26,16 | 53,96 | 98,69 | y = 4,9171x + 1,6744 | 4,92 | 1,00 | 12,27 | 26,16 | 53,96 | 98,69 | 14,12 | 1,00 | 12,27 | 26,16 | 53,96 | 98,69 | 16,75 | 1,00 | 12,27 | 26,16 | 53,96 | 98,69 | 12,81 |

| | Coefficient of variation |
|---|---|
| Slopes not normalised | 65,91 |
| Slopes adjusted at 2,5µg | 3,17 |
| Slopes adjusted at 5µg | 15,19 |
| Slopes adjusted at 10µg | 3,71 |

### Supplementary Table 5

## A

| WB # | Control signal (AU) | | | Equation | Slope | Control signal (AU) | | | Equation | Slope | Internal control signal | Ratio intergel internal | Control signal adjusted by internal control | | | Equation | Slope | OA signal adjusted by internal control | | | Equation | Slope |
|---|---|---|---|---|---|---|---|---|---|---|---|---|---|---|---|---|---|---|---|---|---|---|
| | Protein loaded (µg) | | | | | Protein loaded (µg) | | | | | | | Protein loaded (µg) | | | | | Protein loaded (µg) | | | | |
| | 1 | 5 | 10 | | | 1 | 5 | 10 | | | | | 1 | 5 | 10 | | | 1 | 5 | 10 | | |
| Gel 1 | 1026 | 60762 | 122751 | y= 13479x - 10374 | 13479 | 222357 | 468993 | 818622 | y= 66402x+ 149179 | 66402 | 53656 | 0,27 | 279,48 | 16551,69 | 33437,61 | y = 3671,7x - 2825,9 | 3672 | 60570 | 127755 | 222994 | y = 18088x + 40637 | 18088,00 |
| Gel 2 | 2016 | 12002 | 29193 | y= 3037x- 1793 | 3037 | 43036 | 98112 | 167227 | y= 13800x + 29192 | 13800 | **14616** | 1 | 2016,00 | 12002,00 | 29193,00 | y = 3036,8x - 1792,7 | 3037 | 43036 | 98112 | 167227 | y = 13800x + 29192 | 13800,00 |

## B

| WB # | Control signal (AU)/10^3 | | | Equation | Slope | OA signal (AU) | | | Equation | Slope | Internal control signal | Ratio intergel internal control | Control signal adjusted by internal control | | | Equation | Slope | OA signal adjusted by internal control | | | Equation | Slope |
|---|---|---|---|---|---|---|---|---|---|---|---|---|---|---|---|---|---|---|---|---|---|---|
| | Protein loaded (µg) | | | | | Protein loaded (µg) | | | | | | | Protein loaded (µg) | | | | | Protein loaded (µg) | | | | |
| | 1 | 5 | 10 | | | 1 | 5 | 10 | | | | | 1 | 5 | 10 | | | 1 | 5 | 10 | | |
| Gel 1 | 1,026 | 60,762 | 122,751 | y= 13,479x - 10,374 | 13,479 | 222,357 | 468,993 | 818,622 | y= 66,402x+ 149,179 | 66,402 | 53,656 | 0,27 | 0,28 | 16,55 | 33,44 | y = 3,6717x - 2,8259 | 3,67 | 61 | 128 | 223 | y = 18,088x + 40,637 | 18,09 |
| Gel 2 | 2,016 | 12,002 | 29,193 | y= 3,037x- 1,793 | 3,037 | 43,036 | 98,112 | 167,227 | y= 13,8x + 29,192 | 13,8 | **14,616** | 1,00 | 2,02 | 12,00 | 29,19 | y = 3,0368x - 1,7927 | 3,04 | 43 | 98 | 167 | y = 13,800x + 29,192 | 13,80 |

| | Coefficient of variation |
|---|---|
| Slopes control not normalised | 89,41 |
| Slopes control adjusted by internal control Gel 2 | 13,38 |
| Slopes OA not normalised | 92,75 |
| Slopes OA adjusted by internal control Gel 2 | 19,02 |

| | Before normalization | | After normalization | |
|---|---|---|---|---|
| | Fold change to its control | Fold change to the other control | Fold change to its control | Fold change to the other control |
| | 4,93 | 1,02 | 4,93 | 5,96 |
| | 4,54 | 21,86 | 4,54 | 3,76 |
| Average fold change | 4,74 | 11,44 | 4,74 | 4,86 |
