## Supplementary figures and images for "Titration-WB: A methodology for accurate quantitative protein determination overcoming reproducibility errors"

### Supplementary Figure 1

#### Supplementary Figure 1

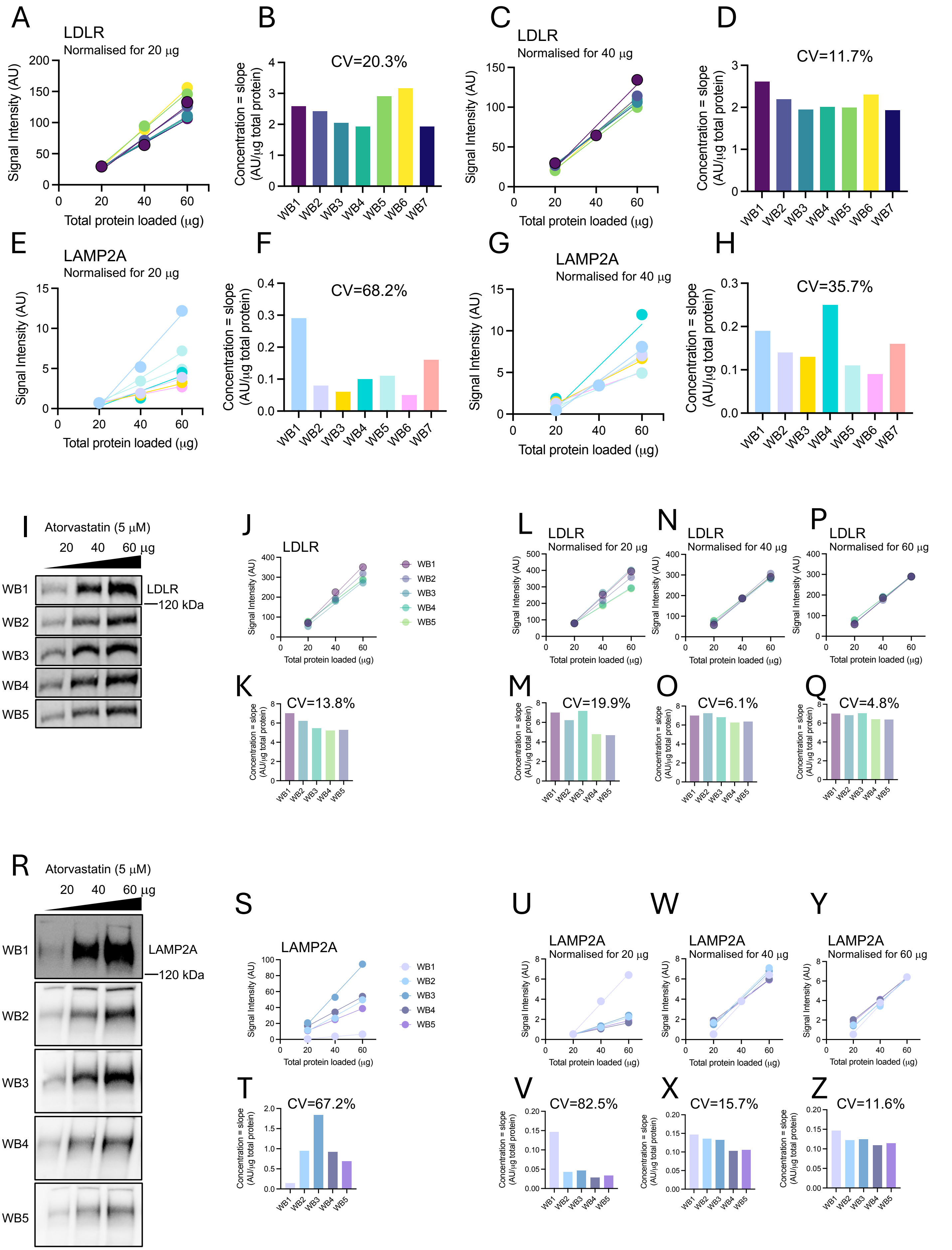
